# Pre-Clinical Application of Functional Human Induced Pluripotent Stem Cell-Derived Airway Epithelial Grafts

**DOI:** 10.1101/2021.05.18.444629

**Authors:** Ratna Varma, Alba E. Marin-Araujo, Sara Rostami, Thomas K. Waddell, Golnaz Karoubi, Siba Haykal

## Abstract

Airway pathologies including cancer, trauma and stenosis lack effective treatments, meanwhile airway transplantation and available tissue engineering approaches fail due to epithelial dysfunction. Autologous progenitors do not meet the clinical need for regeneration due to their insufficient expansion and differentiation, for which human induced pluripotent stem cells (hiPSCs) are promising alternatives. Airway epithelial grafts are engineered by differentiating hiPSC-derived airway progenitors into physiological proportions of ciliated (73.9±5.5%) and goblet (2.1±1.4%) cells on a Silk Fibroin-Collagen Vitrigel Membrane (SF-CVM) composite biomaterial for transplantation in porcine tracheal defects *ex vivo* and *in vivo*. Evaluation of *ex vivo* tracheal repair using hiPSC-derived SF-CVM grafts demonstrate native-like tracheal epithelial metabolism and maintenance of mucociliary epithelium to day 3. *In vivo* studies reveal SF-CVM integration, maintenance of airway patency, showing 80.8±3.6% graft coverage with an hiPSC-derived pseudostratified epithelium and 70.7±2.3% coverage with viable cells, 3 days post-operatively. We demonstrate the utility of bioengineered, hiPSC-derived epithelial grafts for airway repair in a pre-clinical survival model, providing a significant leap for airway reconstruction approaches.

## 1. Introduction

Long-segment airway malignancy, trauma, and stenosis lead to substantial morbidity and mortality. Such advanced cases do not allow for conventional therapeutic approaches, particularly tracheal resection with direct end-to-end anastomosis, resulting in multiple hospitalizations and dependence on permanent tracheostomies and stents.^[1]^ Tracheal transplantation represents a solution for these end-stage disease cases. It is, however, accompanied by many limitations including inadequate revascularization ^[2, 3]^ and immune rejection,^[4]^ but impaired epithelial healing is also a primary cause of transplant failure.^[5]^ Tracheal function is critically dependent on the tracheal epithelium, which moistens the air traveling towards the lungs, acts as a barrier between the internal and external surfaces, and clears the airway of pathogen-entrapping mucous.^[6]^ In fact, the lack of an epithelium itself leads to a fibrogenic response.^[7]^ Therefore, tracheal replacement approaches must meet the clinical need for restoring the pseudostratified airway epithelium.

Tracheal replacements such as stents and protheses have been evaluated clinically, however, such alternatives are limited by complications including infection, stent displacement, inflammation, granulation tissue formation, improper epithelial maturation, and re-stenosis.^[4, 8]^ Decellularized scaffolds ^[9–13]^ and synthetic biomaterials ^[14]^ have demonstrated suboptimal epithelial attachment, survival, migration, and differentiation. These outcomes are particularly appreciated in clinical cases and large animal models. Furthermore, in smaller murine models, epithelial ingrowth from the surrounding tissue can be robust, confounding the assessment of the therapeutic potential of transplanted cells. This has been demonstrated by Okuyama et *al*., where a tissue-engineered tracheal graft was found to be populated with cells originating from the recipient rather than transplanted cells.^[15]^ In general, there has been no compelling evidence of the survival and integration of transplanted cells in tissue-engineered tracheae. To overcome this challenge, focus must be directed towards tracheal tissue engineering approaches that leverage substrates and cell sources promoting epithelial survival and differentiation.

Several naturally derived biomaterials ^[16, 17]^ have demonstrated potential for the development of airway epithelial grafts for surgical use. The absence of defined metrics hinders objective evaluation and comparison of existing and emerging biomaterials in the field. We recently defined a set of quantitative selection criteria for engineering surgically-implantable airway grafts that considered both epithelial support (attachment, metabolic activity, differentiation) as well as biomaterial mechanical properties (bulk degradation and tensile strength). Using these metrics, we systematically compared a wide range of these biomaterials for generating airway epithelial grafts and engineered a composite biomaterial of Silk Fibroin and Collagen Vitrigel Membrane (SF-CVM).^[18]^ This biomaterial provided high tensile strength for surgical manipulation and allowed differentiation of primary human tracheal epithelial cells (HTECs) into ciliated and goblet cells.^[18]^

While primary airway basal cells, such as HTECs, have the potential to differentiate into mature mucociliary epithelium and can be obtained via nasal brushings or bronchoscopy samples, they are limited with respect to their potential for use in tissue engineering applications. Specifically, they are unreliable cell sources due to their high heterogeneity in differentiation capacity across donors, are unable to yield large numbers of cells due to their slow growth rate, and have restricted expansion potential (typically up to two passages), beyond which they lose their ability to differentiate.^[19]^ Moreover, extraction of primary cells from diseased patients may increase variability and limit use in tissue engineering purposes given possible exposure to various environmental and chemical factors.

Human induced pluripotent stem cells (hiPSCs) hold great promise for the development of patient-specific grafts, serving as an expandable cell source, amenable to gene-editing technologies for personalized cell therapy, and addressing the translational limitations of primary cells. Several differentiation protocols exist for generating lung and airway progenitors using pluripotent stem cells, with the primary focus being disease modeling,^[20–26]^ which have been reviewed recently.^[27]^ Only a few studies have evaluated lung and airway cell therapy approaches with hiPSCs,^[15, 28]^ however, these have used murine models and no study to date has been performed in large animal models to establish the clinical utility of hiPSC-derived epithelia.

In this study, we engineered hiPSC-derived airway grafts using our previously established and characterized SF-CVM biomaterial for evaluation in porcine airway defects *ex vivo* and *in vivo*. We achieved desirable differentiation of hiPSC-derived airway progenitors into functional mucociliary epithelium on SF-CVM grafts, which demonstrated maintenance of differentiated epithelia after *ex vivo* airway defect repair and culture. *In vivo*, our hiPSC-derived SF-CVM grafts integrated with the surrounding tracheal tissue, prevented airway obstruction, and maintained a viable, pseudostratified epithelial layer.

## 2. Results

### 2.1 Study Overview

Using an established directed differentiation protocol ^[21]^ (**Figure 1A**), and our previously characterized SF-CVM biomaterial grafts ^[18]^ (**Figure 1B**), our goal was to generate hiPSC-derived SF-CVM airway epithelial grafts (**Figure 1C**) with physiologically relevant proportions of mucociliary epithelial populations (ciliated and goblet cells), and to assess the utility of these grafts both *ex vivo* (evaluating the epithelial differentiation state after airway defect repair) in a bioreactor and *in vivo* (assessing graft integration, airway patency, and epithelial integrity and survival) in a pre-clinical porcine survival model (**Figure 1D**).

**Figure 1.**
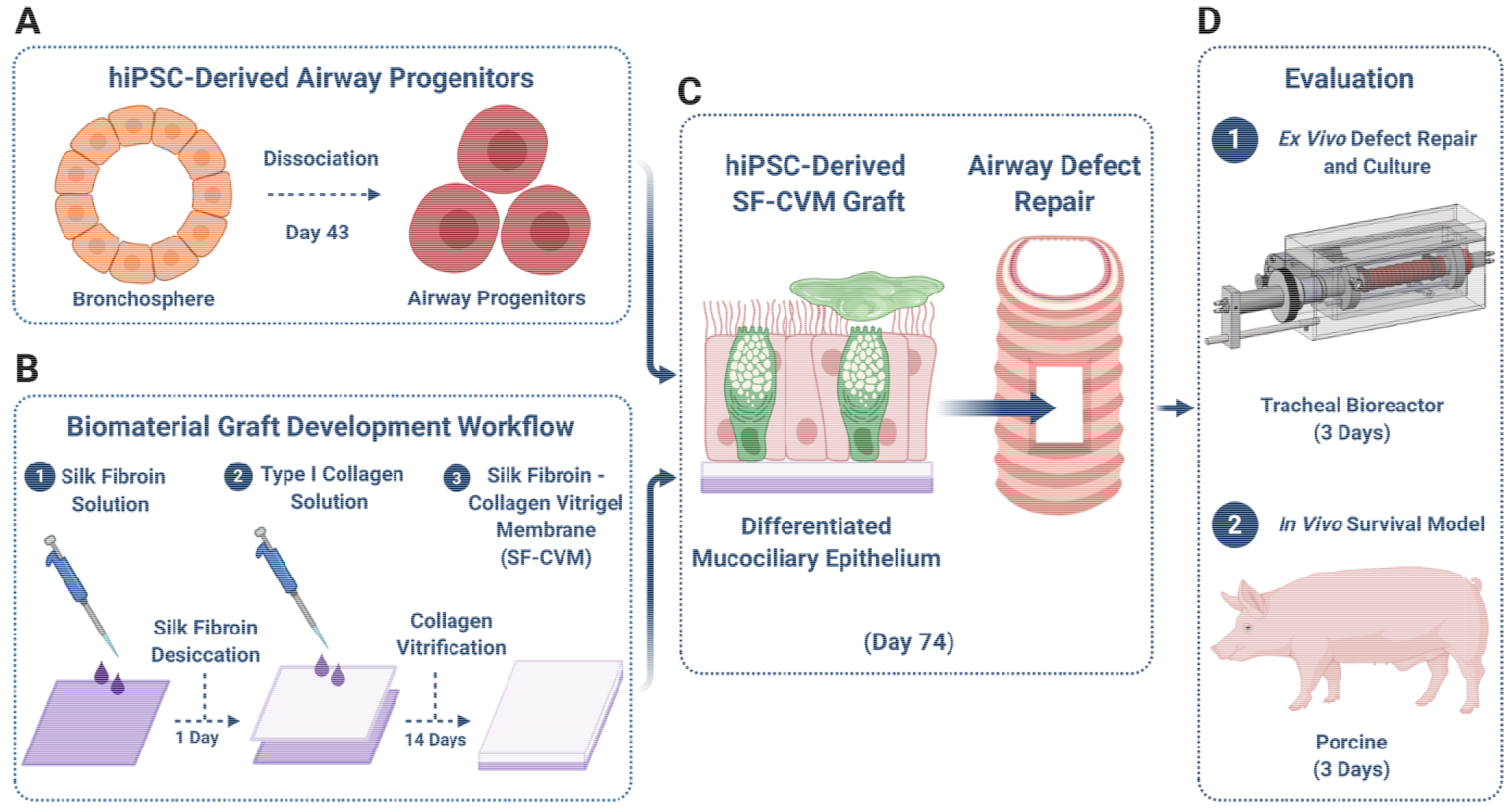
Study Design. Production of hiPSC-derived airway progenitors via directed differentiation (**A**). Workflow of our previously developed composite biomaterial composed of Silk Fibroin and Collagen Vitrigel Membrane (SF-CVM) (**B**). Differentiation of hiPSC-derived airway progenitors on SF-CVM resulting in airway grafts with functional mucociliary epithelia (**C**). Evaluation of hiPSC-derived SF-CVM grafts in *ex vivo* and *in vivo* airway defect models for their airway repair utility (**D**).

### hiPSC-Derived Airway Progenitors Differentiate into Functional Mucociliary Cells on SF-CVM

Through adaptation of an established directed differentiation protocol,^[21]^ we differentiated hiPSCs towards definitive endoderm, lung progenitors, airway progenitors, and mature mucociliary epithelium over 74 days (**Figure 2A**). Using the BU3NG (NKX2.1^GFP^ reporter) line,^[29, 30]^ we produced definitive endoderm populations with 92.5±1.3% CKIT^+^CXCR4^+^ expression (**Figure 2B**). The 110 hiPSC line (**Figure S1A**) similarly resulted in 91.0±0.7% CKIT^+^CXCR4^+^ definitive endoderm on day 3 (**Figure S1B**). By day 15, we induced lung progenitor populations using the BU3NG line with 57.1±10.1% of cells expressing NKX2.1^GFP^ (**Figure 2C**). We confirmed the validity of this reporter through intracellular staining for NKX2.1 which showed a high degree of concordance (**Figure 2C**). We achieved a similar induction of lung progenitors using the 110 hiPSC line, with 33.6±4.1% of cells demonstrating CD47^hi^CD26^lo^ expression, a phenotype correlated with NKX2.1 expression,^[29]^ by day 15 (**Figure S1C**) which was comparable to that of BU3NG lung progenitors (*P*=0.10; **Figure S1D**).

**Figure 2.**
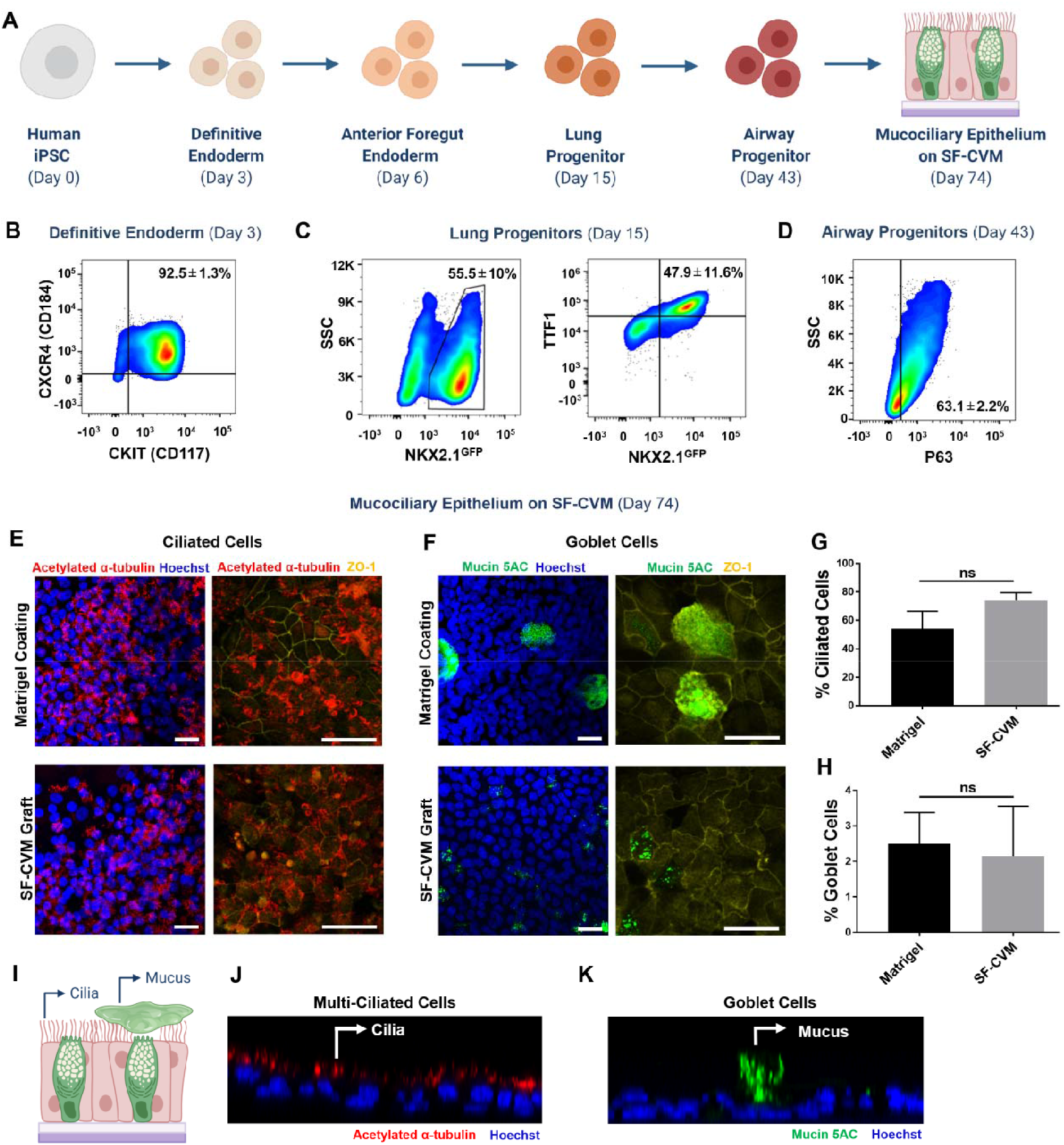
Functional mucociliary SF-CVM airway epithelial grafts derived via directed differentiation of hiPSCs. Directed differentiation schematic for derivation of mucociliary airway epithelium on SF-CVM (**A**). Representative flow cytometry dot plot analysis of CKIT^+^CXCR4^+^ cells of day 3 definitive endoderm (**B**), NKX2.1^GFP+^ and NKX2.1^GFP+^ TTF1^+^ cells of day 15 lung progenitors (**C**), and P63^+^ cells of day 43 airway progenitors (**D**). Differentiation comparison of hiPSC-derived airway progenitors into ciliated **(E, G)** and goblet **(F, H)** cells under ALI culture for 31 days (day 74) on Matrigel-coated (standard control) and SF-CVM-laden transwell inserts with no significant differences observed between ciliated **(G)** and goblet **(H)** cell populations. Cross-sectional schematic of functional mucociliary epithelium on SF-CVM **(I)**. Presence of multi-ciliated **(J)** and mucus secretion above the epithelial layer by goblet cells **(K)** on hiPSC-derived SF-CVM grafts. Groups of two were analyzed via two-tailed Student’s t-test. (Mean ± SEM; *N*=3; Scale bars: 25 μm)

Lung progenitor populations were sorted and embedded in Matrigel to form bronchospheres with both the BU3NG line (**Figure S1E**) and 110 line **(Figure S1F)**. Dissociated BU3NG bronchosphere cells demonstrated 63.1±2.2% P63^+^ expression (**Figure 2D**) by day 43. These day 43 airway progenitors were seeded on either Matrigel-coated (standard control) or SF-CVM-laden transwell inserts and subsequently maintained in air-liquid interface (ALI) culture to promote differentiation towards ciliated and goblet cells by day 74. BU3NG hiPSC-derived airway progenitors (hiPSC-APs) differentiated into 54.2±12.2% and 73.9±5.5% ciliated cells (**Figure 2E,G**) on Matrigel and SF-CVM, respectively, with no significant differences found between the two populations (*P*=0.21; **Figure 2G**). The percentage of ciliated cells produced by BU3NG hiPSC-APs was significantly higher than that of HTECs (23.7±8.5%) on SF-CVM (*P*<0.05; **Figure S2A,B**), indicating superior performance of hiPSC-APs in inducing ciliary differentiation than primary HTECs. Meanwhile, differentiation of BU3NG hiPSC-APs into goblet cells resulted in 2.5±0.9% and 2.1±1.4% goblet cells (**Figure 2F,H**) on Matrigel and SF-CVM, respectively, with no significant differences found (*P*=0.85; **Figure 2H**). There were also no significant differences between differentiation of BU3NG hiPSC-APs and HTECs (6.6±2.8%) into goblet cells on SF-CVM (*P*=0.39; **Figure S2C,D**). Additionally, the BU3NG hiPSC-derived epithelia on SF-CVM demonstrated consistent ZO-1 expression throughout the grafts, indicating epithelial barrier integrity (**Figure 2E,F**). Notably, 110 hiPSC-APs lost their epithelial monolayer on Matrigel-coated transwells, resulting in a patchy epithelium (**Figure S1G**). On SF-CVM, however, 110 hiPSC-APs resulted in a uniform epithelial layer with 40.1±16.2% ciliated cell (*P=*0.12; **Figure S1H,I**) and 2.6± 2.1% goblet cell (*P=*0.87; **Figure S1J,K**) populations, which were comparable to those produced by BU3NG hiPSC-APs. Although ciliated cell populations of both hiPSC lines on SF-CVM were statistically similar, those of the 110 line were immature and single-ciliated with minimal presence of multi-ciliated cells by day 74. In contrast, BU3NG hiPSC line primarily led to multi-ciliated cell populations on SF-CVM (**Figure 2E**). Prior to moving forward with the BU3NG line for subsequent experiments, we confirmed the differentiation quality of its derived mucociliary epithelium on SF-CVM by assessing epithelial cross-sections for cilia projections and mucus secretion by goblet cells (**Figure 2I**). We observed multiple mature ciliary projections (**Figure 2J**), as well as evidence of functional mucus secretion above the epithelial layer by goblet cells (**Figure 2K**).

Previous studies have demonstrated increased ciliated cell differentiation of primary or hiPSC-derived airway progenitors via introduction of basic fibroblast growth factor (bFGF),^[17]^ insulin, ^[31]^ interleukin-6 (IL-6),^[32]^ WNT signaling,^[33]^ and Notch inhibition ^[22, 25, 34]^ during ALI culture. We attempted to enhance ciliation of BU3NG hiPSC-APs on SF-CVM by subjecting them to bFGF, DAPT, DBZ, dickkopf-1 (DKK-1), insulin growth factor-1 (IGF-1), and IL-6 during ALI culture. None of the groups performed better than SF-CVM alone with regards to ciliation, and there were no statistical differences amongst all groups for both ciliated (**Figure S3A**) and goblet (**Figure S3B**) cell differentiation. Given that SF-CVM was able to independently induce physiological proportions of ciliated cells, all subsequent experiments were performed on BU3NG hiPSC-derived SF-CVM grafts without additional modifications.

### 2.3 hiPSC-Derived SF-CVM Airway Grafts Maintain Their Differentiated State in *Ex Vivo* Airway Defects

We modified our previously described perfusion-based bioreactor ^[35, 36]^ system to allow for ALI culture (**Figure S4A**). To assess the impact of ALI culture, we compared tracheal epithelial histology of 5 cm-long native pig tracheae cultured in the submerged (**Figure S4B**) or ALI (**Figure S4C**) culture conditions for 7 days. On day 7, in submerged culture, the tracheal epithelium was devoid of ciliated cells and had reached a squamous state throughout the length of the trachea (**Figure S4D**), while a pseudostratified, ciliated epithelium was maintained throughout the tracheal length in ALI culture (**Figure S4E**).

Using our bioreactor platform, we examined the behavior of our differentiated SF-CVM grafts in porcine airway defects under *ex vivo* ALI culture conditions. We created a 4 cm x 2 cm defect in the trachea already set up in the bioreactor, manually stripped the mucosa of the defect piece, attached the hiPSC-derived SF-CVM airway graft, and closed the defect (**Figure 3A**). We evaluated the epithelial metabolic activity of native tracheae and tracheae with defect repair using our hiPSC-derived SF-CVM grafts by measuring the bicarbonate (**Figure S4F**), glucose (**Figure S4G**), and lactate (**Figure S4H**) concentrations in collected media samples across the three days of *ex vivo* ALI culture. Metabolite measurements for both tracheal groups were statistically similar across the three days. We further evaluated the hiPSC-derived mucociliary epithelial state on these grafts before (**Figure 3B**) and after (**Figure 3C**) defect repair and *ex vivo* ALI culture, finding that both ciliated and goblet cell phenotypes were maintained across the three days.

**Figure 3.**
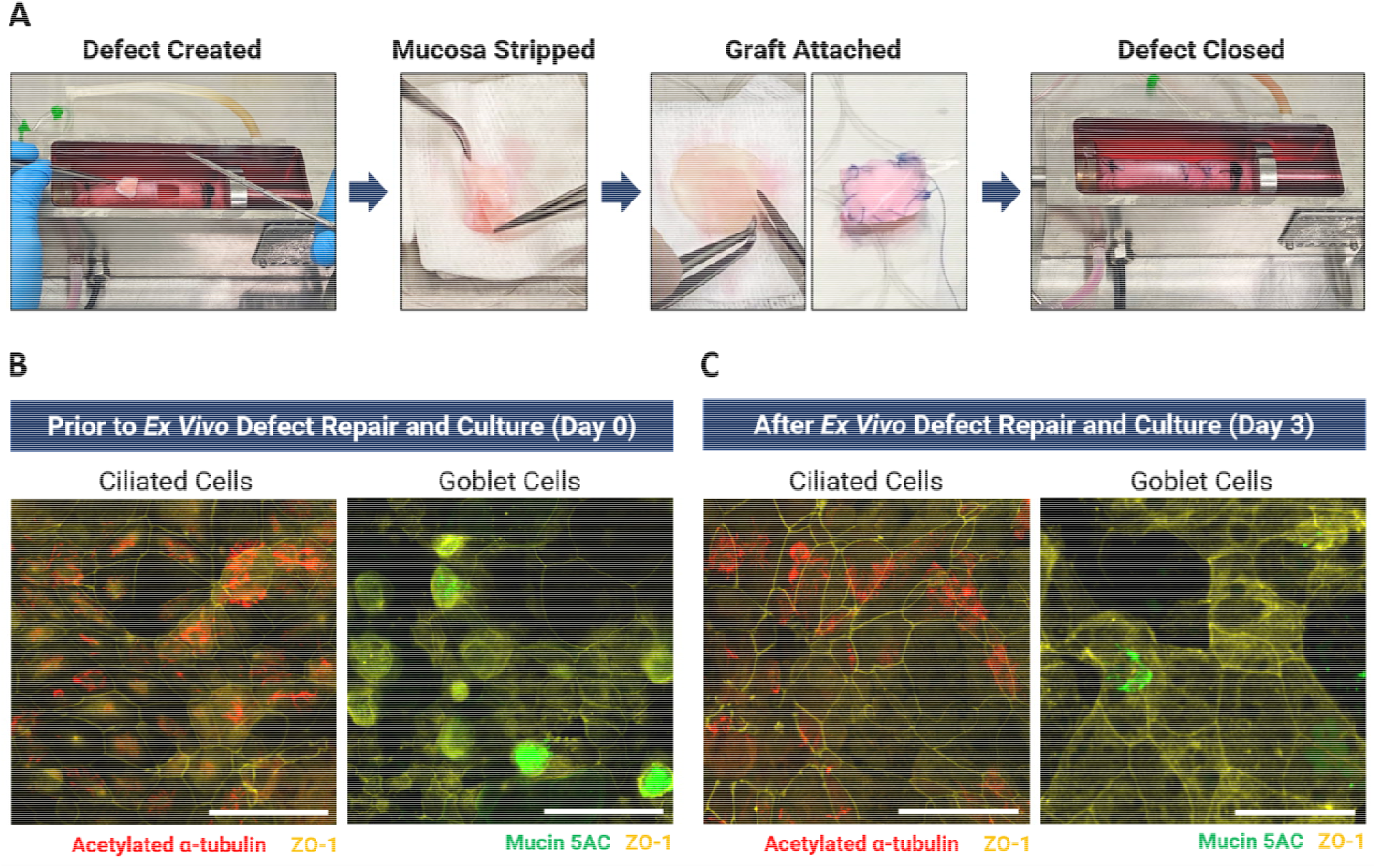
hiPSC-derived SF-CVM airway grafts maintain mucociliary epithelium after 3 days of *ex vivo* ALI culture. Workflow of porcine airway defect creation and repair using hiPSC-derived SF-CVM airway grafts within a bioreactor (**A**). After defect closure, hiPSC-derived SF-CVM airway grafts were cultured for 3 days in a double chamber perfusion-based bioreactor with a peristaltic unidirectional flow at a rate of 1.5 mL min^−1^, an inner chamber rotational speed of 1 rpm, and simultaneous injection of air and media into the inner chamber for ALI culture. Differentiated ciliated and goblet cell phenotypes are shown on SF-CVM grafts prior to (**B**) and after (**C**) 3 days of *ex vivo* ALI culture. *N*=3; Scale bars: 25 μm.

### 2.4 hiPSC-Derived SF-CVM Airway Grafts Integrate with Surrounding Tissue, Maintain Their Pseudostratified Epithelial Integrity and Viability, and Reduce Host Inflammatory Response Following *In Vivo* Transplantation

After confirming successful *ex vivo* culture of hiPSC-derived SF-CVM grafts within airway defects, we developed a porcine airway defect model for *in vivo* studies of graft integration and survival across 3 days. Herein, we created a 4 cm x 2 cm tracheal defect and manipulated it according to the following four groups prior to defect closure: 1) control (no manipulation); 2) mucosa stripped; 3) mucosa stripped and replaced with bare SF-CVM graft; and 4) mucosa stripped and replaced with hiPSC-derived SF-CVM graft (**Figure 4A**). All animals survived without respiratory or other health complications. On post-operative day (POD) 3, the control group demonstrated near-complete mucosal healing at the site of defect (**Figure 4B**) and a patent airway (**Figure 4C**) macroscopically. Histological analysis of the defect area revealed a thick pseudostratified epithelium (**Figure 4D**), indicating possible hyperplasia, and a lack of ciliated cells in contrast to native tracheal epithelium (**Figure 4E**). The stripped mucosa group showed macroscopic evidence of airway obstruction, which was consistent throughout the longitudinal length of the defect (**Figure 4B, C**). Histologically, the defect region was devoid of a pseudostratified, ciliated epithelium (**Figure 4D**). Both groups with SF-CVM grafts, either bare or with hiPSC-derived epithelium, demonstrated SF-CVM graft integration with the surrounding tracheal tissue with hyperplasia along the edges of the defects (**Figure 4B**), as well as unobstructed airways (**Figure 4C**) macroscopically. Bare SF-CVM grafts remained intact on POD 3, however no evidence of an epithelial layer was observed on their surfaces (**Figure 4D**). Meanwhile, a pseudostratified epithelial layer was evident on the SF-CVM biomaterial surface, albeit demonstrating a lack of ciliated cells (**Figure 4D**), similar to the control group. To evaluate the origin of the pseudostratified epithelium, we tracked the hiPSC-derived epithelium before and after transplantation using a cell tracker and further assessed cell viability of all groups (**Figure 5A**). Cells on hiPSC-derived SF-CVM grafts were labeled with 7-amino-4-chloromethylcoumarin (CMAC) dye and the majority of cells was also positive for calcein-AM (live cell marker) prior to airway defect surgery on POD 0 (**Figure 5B**). On POD 3, 80.8±3.6% of the area of the SF-CVM graft was covered with CMAC-labeled cells (**Figure 5C**). At the same time, 70.7±2.3% of the SF-CVM graft area was covered with calcein-AM^+^ cells (**Figure 5C**). Although we did not stain specifically for native porcine cells, the alignment of CMAC-negative and calcein-AM-negative areas was consistent with approximately 20% of the graft being devoid of any cells whatsoever. Evaluation of the epithelial differentiation state on POD 3 (**Figure 5A**) of hiPSC-derived SF-CVM grafts revealed an epithelial layer with abnormal zonula occludens-1 (ZO-1) localization, a loss of ciliated cells with minimal evidence of single cilia-like punctate structures (**Figure 5E**), and maintenance of mucin 5AC expression (**Figure 5F**). Importantly, cross-sectional imaging of freshly extracted POD 3 grafts demonstrated a viable, pseudostratified, and hiPSC-derived epithelium on SF-CVM (**Figure 5G**).

**Figure 4.**
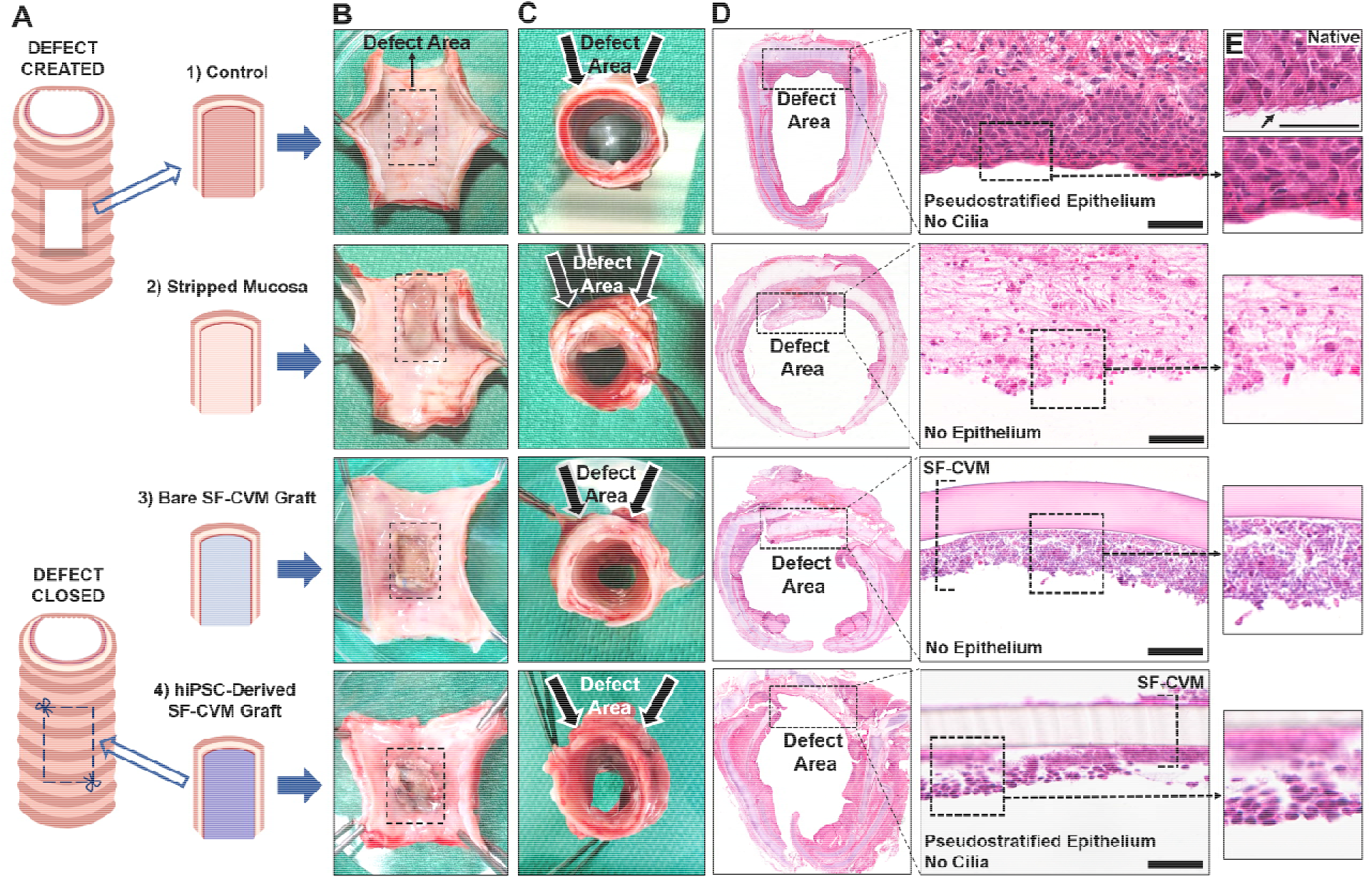
hiPSC-derived SF-CVM grafts integrate with tracheal tissue, demonstrate airway patency and maintain a pseudostratified epithelial layer. Schematic of the *in vivo* porcine airway defect model (**A**). Macroscopic images of exposed tracheal luminal surfaces (**B**) and top (proximal) views of tracheal tubes (**C**) for all experimental groups. Hematoxylin and eosin staining of tracheal cross-sections with insets depicting presence (groups 1 and 4) or absence (groups 2 and 3) of a pseudostratified epithelium within defect areas (**D**). Reference of native tracheal epithelium with cilia (indicated by arrow) (**E**). *N*=3 (Groups 1-3), *N*=4 (Group 4); Scale bars: 50 μm.

**Figure 5.**
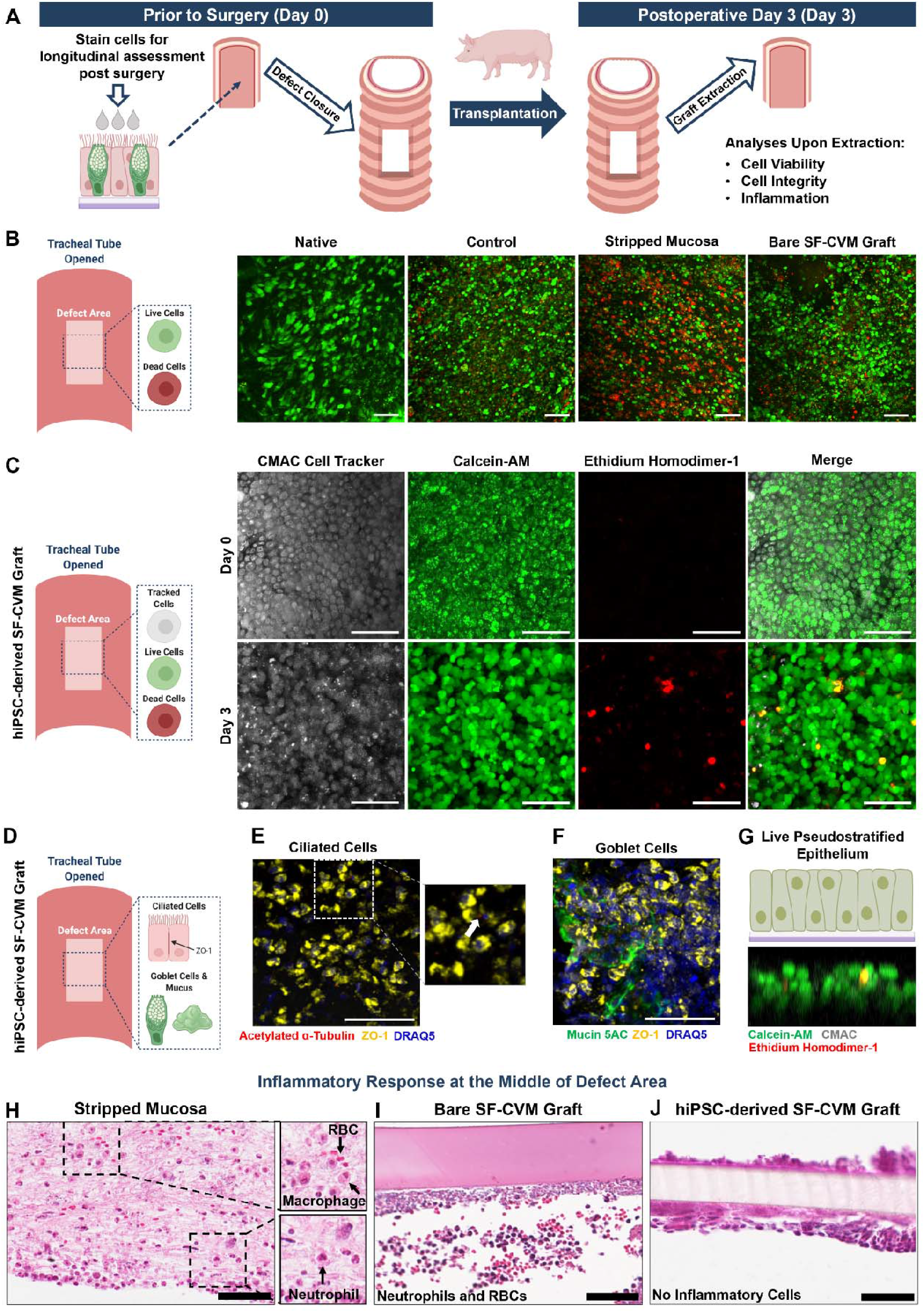
hiPSC-derived SF-CVM grafts maintain epithelial integrity and viability, and reduce inflammation by POD 3. Schematic of hiPSC-derived SF-CVM graft preparation prior to surgery and evaluation post transplantation (**A**). Cell viability assessment of native tracheal epithelium and defect regions of control, stripped mucosa, and bare SF-CVM graft groups on postoperative (POD) 3 (**B**). Images portraying intact and viable epithelial layer of hiPSC-derived SF-CVM grafts tracked prior to surgery (Day 0) and on POD 3 (Day 3) (**C**). Evaluation of epithelial differentiation (**D**) with immunofluorescence images depicting punctate single cilia structure (**E;** indicated by arrow) and presence of mucus (**F**). Cross-sectional view of a live, pseudostratified hiPSC-derived epithelium (**G**). Hematoxylin and eosin staining of defect regions corresponding to stripped mucosa (**H**), bare SF-CVM graft (**I**), and hiPSC-derived SF-CVM graft (**J**) groups on POD 3, depicting no infiltration of inflammatory cells (macrophages, neutrophils, and red blood cells (RBC) on hiPSC-derived SF-CVM grafts. Scale bars: 50 μm.

We further evaluated the host inflammatory response at the site of defect through histology. In the stripped mucosa group, which developed macroscopical airway obstruction, the entire defect region was infiltrated with inflammatory cells including macrophages, neutrophils, and red blood cells (RBCs) trapped within a fibrous matrix (**Figure 5H**). Lesser infiltration of neutrophils and RBCs was observed on the surface of bare SF-CVM grafts (**Figure 5I**), while there was limited evidence of an inflammatory response in defects repaired with hiPSC-derived SF-CVM grafts (**Figure 5J**).

## 3. Discussion

Functional epithelia derived from human pluripotent cell sources hold immense promise for application in lung and airway regeneration therapies. In this study, we developed an hiPSC-derived airway epithelial graft, based on a composite SF-CVM biomaterial, which contained physiologically relevant numbers of ciliated and goblet cells. These grafts maintained their integrity and differentiated state of cells within a porcine airway defect across 3 days of *ex vivo* ALI culture in a bioreactor. *In vivo* repair of porcine airway defects with hiPSC-derived SF-CVM grafts was successful such that no respiratory complications or mortality were experienced by the animals, the grafts integrated well with surrounding tracheal tissue as well as maintained airway patency. Furthermore, the vast majority of the cells on SF-CVM at the site of defect were hiPSC-derived and viable on POD 3, depicting a pseudostratified epithelium.

Moving towards personalized medicine necessitates development of grafts with autologous cells. Particularly, use of recipient-derived cells can minimize the need for immunosuppressive therapy, which otherwise increases susceptibility for infections and complications that in turn decrease life expectancy of patients.^[4]^ Our results demonstrated that hiPSC-APs lead to significantly higher production of ciliated cells compared to primary HTECs, thereby bolstering the clinical relevance of hiPSC sources. Since the emergence of various directed differentiation protocols for lung and airway epithelia, the major focus has been the application of these protocols for disease modeling.^[27]^ As such, little attention has been given to developing epithelial populations that emulate proportions present in the human airway (48-70% ciliated cells ^[37, 38]^ and 6-25% goblet cells ^[38, 39]^). Generating physiological proportions of airway specific cell types is necessary to enable the use of hiPSC-derived epithelial cells in cell-based tissue engineering applications. The proportion of ciliated cells, measured through protein expression of Forkhead box protein J1 (FOXJ1) or acetylated tubulin, has been quantified by Wong et *al*.,^[23]^ de Carvalho et *al*., ^[24]^ and Dye et *al*. ^[26]^ as being 36%, 14%, and 3%, respectively. Meanwhile, the mucin 5B expressing goblet cell population was found to be less than 1% by de Carvalho et *al*. ^[24]^ and mucin 5AC expressing cells was reported as 1-2% by Firth et *al*. ^[22]^. In contrast to our results, the aforementioned protocols developed substantially lower ciliated cell populations than what is considered physiologically relevant. Beyond production of functional cell types from hiPSCs, evaluating the therapeutic potential of these cells following orthotopic transplantation is essential. In our case, we demonstrated presence of a pseudostratified epithelium on SF-CVM, wherein 80.8±3.6% of the SF-CVM graft area was covered with hiPSC-derived cells and a similar proportion of the graft (70.7±2.3%) was covered with live cells on POD 3.

Although the majority of the hiPSC-derived epithelium on SF-CVM was retained at POD 3, abnormal Zonula occludens-1 staining suggests that there was some disruption of the epithelial barrier function. This was accompanied by an additional loss of ciliated cells, which was mirrored by the control group. This epithelial state has been previously described by Inayama et *al*., who resected segments of rabbit tracheae and re-anastomosed them to observe epithelial morphology across time.^[40]^ They observed a ciliated, pseudostratified columnar epithelium on POD 1, which degraded to a columnar epithelium with severe loss of ciliated cells by POD 4. While the epithelium regained its pseudostratified morphology by POD 7, ciliated cells did not appear until POD 10. Murine models of tracheal transplant have also exhibited this pattern of epithelial repair,^[41]^ supporting our POD 3 observations.

Although our SF-CVM grafts and survival model demonstrated promise for tracheal tissue engineering, there were some limitations. While induction of ciliated cells on our hiPSC-derived SF-CVM grafts was well within the physiological range, that of goblet cells was lower, however, goblet cells may be increased via Notch agonism during ALI culture at the cost of decreasing ciliated cell numbers.^[34]^ We also encountered variable differentiation across different hiPSC cell lines, which was likely a function of inherent genetic differences amongst hiPSC donors,^[42]^ similar to those seen across primary cell populations. This variability can be addressed through further refinements of the directed differentiation protocol as well as optimization of the protocol for each hiPSC donor. Another limitation of our survival model was its short time frame, however, we expect that a long-term survival model will exhibit an epithelial repair chronology similar to that described in the aforementioned Inayama et *al*. study, and allow re-establishment of the ciliated, pseudostratified morphology of the hiPSC-derived epithelium. Moreover, given the available pig species and cellular products, immune suppression would need to be administered, potentially confounding interpretation of the results.

A frequent complication of synthetic or tissue-engineered tracheal grafts is overgrowth of granulation tissue and airway narrowing.^[2, 12]^ In this study, we observed prominent presence of non-epithelial cell types in the stripped mucosa group which included macrophages, neutrophils, and red blood cells entrapped in a fibrous matrix. The combination of these factors is indicative of granulation tissue, which is essential for wound healing.^[43]^ In cases of excessive, or chronic injury, however, overgrowth of granulation tissue can occur, which entails excessive deposition of extracellular matrix, forming scar tissue, as well as airway narrowing.^[44]^ Initiation of this process was seen in the stripped mucosa group, especially with the macroscopic assessment displaying airway narrowing. In contrast, we observed minimal cell infiltration, which was primarily present at the edges of the defect when SF-CVM grafts (with or without cells) were implanted. This infiltration only existed in places where the SF-CVM was perforated around sutures, as seen in the macroscopic images as well as in the middle of the defect region on the surface of bare SF-CVM grafts. While the biomaterial itself contributed as a barrier to minimize initiation of granulation tissue formation, as seen in the stripped mucosa group, presence of the hiPSC-derived pseudostratified epithelium on SF-CVM appeared to reduce the inflammatory response. This finding is in congruence with earlier studies which have established the role of an epithelial barrier in preventing inflammation.^[7]^

Another advantage of our hiPSC-derived SF-CVM grafts is their generation *in vitro*, which only requires a 5×10^5^ cells (cm^2^)^−1^ seeding density of airway progenitors. This seeding density is 10-50% of that previously reported for tracheal tissue engineering approaches with primary airway progenitors,^[13, 45]^ making it a feasible option for generating larger grafts in the future. These are all important characteristics of the SF-CVM graft as it requires lesser cell production for seeding, does not impede the initial healing process, facilitates migration of host cells (as seen in the bare SF-CVM group), and maintains the implanted hiPSC-derived pseudostratified epithelium.

Large animal models such as pigs, sheep, dogs, and goats provide an advantage for *in vivo* studies as their airway size, anatomy, and physiology are comparable to those of humans.^[47]^ Based on this, our porcine airway defect model is predictive of hiPSC-derived airway graft behavior in physiological airway environments, and therefore provides a significant leap in application of tissue engineering approaches for the trachea. Additionally, in contrast to heterotopic transplantations of tissue-engineered tracheal grafts, our orthotopic defect model is a high-risk intervention, especially given any airway complications can severely impact animal health, possibly leading to death. Lack of animal mortality and airway concerns throughout our study speak to the success of these hiPSC-derived SF-CVM airway grafts.

Overall, our hiPSC-derived SF-CVM grafts demonstrate immense clinical relevance based on their ability to induce mature mucociliary epithelia. Their ability to maintain viable respiratory epithelium as well as preserve airway patency is especially promising. Not only do these grafts show promise for tracheal reconstruction, but also serve as a stepping-stone for generating and evaluating epithelial grafts for other tubular organs.

## 4. Conclusion

Tissue-engineered hiPSC-derived SF-CVM airway grafts were developed with physiological proportions of differentiated mucociliary epithelial cells. *Ex vivo* porcine airway defect repair using these grafts, within a bioreactor, revealed maintenance of respiratory epithelium, while *in vivo* airway defect repair in porcine survival model demonstrated graft integration without airway obstruction as well as presence of a viable, pseudostratified hiPSC-derived airway epithelium. Our translational approach shows promise for the long-term adaptation of hiPSC-derived grafts within complex pre-clinical models, thus providing a significant advancement for the field of airway reconstruction.

## 5. Experimental Section/Methods

### 5.1 Ethical Considerations

Human induced pluripotent stem cell lines were approved in accordance with the guidelines provided by the Stem Cell Oversight Committee of the Canadian Institute of Health Research. All animals in this study received humane care as outlined in the “Principles of Laboratory Animal Care” provided by the National Society for Medical Research and the “Guide for the Care of Laboratory Animals” published by the National Institutes of Health. All studies were approved by the Animal Resource Centre of the Toronto General Hospital Research Institute.

### 5.2 hiPSC Maintenance and Differentiation

BU3NG NKX2.1^GFP^ reporter hiPSC line (a generous gift from Dr. Darrell Kotton, Boston University) was cultured in feeder-free conditions on Matrigel-coated (hESC-qualified; Corning) plates in mTESR1 media (Stem Cell Technologies) and passaged as small clumps using Gentle Cell Dissociation Reagent (GCDR; Stem Cell Technologies) for 3 minutes. The 110 hiPSC line (a kind gift from Dr. Andras Nagy, University of Toronto) was cultured in feeder-free conditions on Geltrex®-coated (Thermo Fisher) dishes in mTESR1 media (Stem Cell Technologies) and passaged as single cells using TrypLE™ Express Enzyme (Thermo Fisher) for 3 minutes. Lung and airway progeny were generated as previously described.^[21]^ Unless otherwise stated, all differentiation was performed in serum-free differentiation media (SFDM) containing Iscove’s Modified Dulbecco’s Medium (Thermo Fisher) and Ham’s F-12 (Thermo Fisher) at a 3:1 ratio with 0.5% N2 and 1% B27 supplements (Thermo Fisher), 1% GlutaMax (Thermo Fisher), 1% penicillin/streptomycin (Wisent), 0.05% bovine serum albumin (Thermo Fisher). Complete SFDM (cSFDM) was made via fresh supplementation with 50 μg mL^−1^ ascorbic acid (Sigma-Aldrich) and 0.4 μм monothioglycerol (Sigma-Aldrich) at the time of use.

Definitive endoderm was induced using the STEMdiff^TM^ Definitive Endoderm Kit (Stem Cell Technologies) with a slight modification to the manufacturer’s protocol, wherein cells were harvested at 72 hours (day 3), instead of 96 hours for flow cytometry analysis or further differentiation. These cells were dissociated as clumps using GCDR for 3 minutes, seeded onto Matrigel-coated (hESC-qualified; Corning) plates, and cultured in cSFDM with 2 μм Dorsomorphin (Tocris) and 10 μм SB431542 for 72 hours (day 3-6) for patterning anterior foregut endoderm. Subsequently, these cells were subjected to cSFDM with 3 μм CHIR99021 (Tocris), 10 ng mL^−1^ recombinant human bone morphogenic protein (BMP4; R&D Systems), and 50 nм retinoic acid (RA; Sigma-Aldrich) between day 6 and 15 for production of lung progenitors. Between days 14 and 16, cells were dissociated for 20 minutes using 0.5% Trypsin-EDTA (Thermo Fisher) and sorted for NKX2.1-GFP expression (BU3NG line; *N*=4) or CD47^hi^ CD26^lo^ expression (110 line; *N*=4) ^[29]^. Sorted cells were embedded in 100% Matrigel (growth-factor reduced; Corning) drops at 1000 cells μL^−1^ and cultured in cSFDM with 250 ng mL^−1^ fibroblast growth factor 2 (FGF 2; R&D Systems), 10 ng mL^−1^ FGF10 (R&D Systems), 10 μм Y27632 (Tocris), 50 nм dexamethasone (Sigma), 0.1 mM 8-Bromoadenosine 30, 50-cyclic monophosphate sodium salt (Sigma) and 0.1 mм 3-Isobutyl-1-methylxanthine (Sigma) for 4 weeks to generate airway progenitor organoids by Day 41 to 43. Airway progenitors were dissociated into single cells using 2 mg mL^−1^ Dispase II (Sigma Aldrich) for 1 hour, followed by 20 minutes of 0.5% Trypsin-EDTA incubation for flow cytometry analysis or further differentiation to generate airway epithelial grafts.

### 5.3 Generation of hiPSC-derived SF-CVM Airway Epithelial Grafts

SF-CVM bilayer (2.5 cm disc) was fabricated as previously described.^[18]^ Hydrated SF-CVM discs were dabbed on gauze, with the SF side facing the gauze, to remove excess Dulbecco’s Phosphate-Buffered Saline (DPBS; Thermo Fisher) and glued onto 6-well transwell inserts (Corning) using Fibrin Glue (Baxter International). SF-CVM was allowed to bond with the transwell and simultaneously sterilized under UV radiation in a sterile biosafety cabinet overnight. SF-CVM was rehydrated with DPBS in the transwells prior to cell seeding. Day 41-43 airway progenitors were seeded on SF-CVM at 500,000 cells (cm^2^)^−1^ and allowed to form a monolayer in submerged culture for 5 days in complete PneumaCult-ALI^TM^ (Stem Cell Technologies) media (containing dexamethasone instead of hydrocortisone, 1% Antibiotic-Antimycotic (Thermo Fisher), and 1% gentamicin (Wisent)) with further supplementation of 2 μм Dorsomorphin and 10 μм SB431542. On day 5, air-liquid interface (ALI) culture was begun by adding, to the bottom chamber only, complete PneumaCult-ALI^TM^ media for an additional 31 days (day 72-74). The resulting grafts were assessed for terminal differentiation via immunocytochemistry and/or used for *ex vivo* and *in vivo* studies.

### 5.4 *Ex Vivo* Studies

#### 5.4.1 Tracheal Collection and Decontamination

Whole cadaveric tracheae from outbred male Yorkshire donor pigs (30-40 kg) were procured in a sterile fashion using standard surgical procedures.^[10, 13, 35]^ The tracheal lumen was scraped with a metal spatula to remove residual mucous and washed on a shaker for 1 hour at room temperature in cold decontamination solution containing Dulbecco’s Modified Eagle Media (DMEM; Thermo Fisher) with 10% fetal bovine serum, 2 mg mL^−1^ fluconazole, 25 mg mL^−1^ colistimethate, 67.5 mg mL^−1^ imipenem/cilastatin, 385 mg mL^−1^ ceftazidime, 2% Antibiotic-Antimycotic (Thermo Fisher), 0.1% Primocin^TM^ (InvivoGen), and 0.1% Fungin^TM^ (InvivoGen). Subsequently, the scraping and washing steps were repeated twice more, resulting in a total decontamination wash period of 3 hours.

#### 5.4.2 Bioreactor Culture System

A double chamber perfusion bioreactor, developed by our laboratory, was assembled as previously described ^[13, 35, 36]^ in a sterile biosafety cabinet. Peristaltic unidirectional flow at a rate of 1.5 mL min^−1^, and rotation of inner chamber of 1 rpm (luminal wall shear stress 148 μPa) were utilized. The outer chamber of the bioreactor was filled with cartilage-supportive media as previously described. For submerged culture experiments, the tracheal luminal chamber was filled with PneumaCult-Ex^TM^ media. For ALI experiments, the tracheal lumen was purged of all media, following which filtered humidified air and media were pumped into the trachea with the bioreactor positioned at a 30° inclination. A dripping system was created to simultaneously expose the epithelium to air and intermittent droplets of PneumaCult-ALI^TM^ media. Native pig tracheae (*N*=3) were cultured in submerged or ALI culture for 7 days in the bioreactor and subsequently assessed via histology.

#### 5.4.3 Ex Vivo Airway Defect Study

Decontaminated tracheae were set up in the bioreactor for ALI culture in sterile conditions. A 4 cm x 2 cm tracheal defect was created near the inlet of the bioreactor, and the piece was removed for manipulation. The pig mucosa surface including the epithelium was manually stripped using forceps. The remaining graft was briefly dabbed with gauze and Fibrin Glue (Baxter International) was applied. The hiPSC-derived SF-CVM graft was cut to match the defect shape and further secured onto the defect via 4-0 PROLENE® polypropylene running sutures. This segment was subsequently re-attached to the trachea via 4-0 PROLENE® polypropylene sutures to close the defect with the cells being oriented towards the luminal surface. The tracheae were cultured in ALI for 3 days. Media samples were collected daily from the tracheal luminal media circuit to evaluate metabolite concentrations (bicarbonate, glucose, and lactate) across *ex vivo* ALI culture. Samples were analyzed using RAPIDPoint 500 Blood Gas Systems (Siemens Healthcare Limited). After three days of ex vivo culture, hiPSC-derived SF-CVM grafts were extracted to assess the epithelial differentiation state via immunocytochemistry.

### 5.5 *In Vivo* Airway Defect Study

#### 5.5.1 Animal Husbandry and Monitoring

Outbred Yorkshire male pigs (35-40 kg) were used for the studies. All animals were monitored for any signs of severe illness, including diarrhea, respiratory distress, lameness, and severe skin lesions, and acclimated for at least 72 hours prior to experimental procedures. Prior to and post experimental procedures, all animals were housed in 2 × 4 square meter pens with the following environmental conditions: 10-25°C temperature and 55-75% relative humidity. The pigs were provided water *ad libitum* via a sipper and fed according to the 8753 Teklad (Harlan) Miniswine Diet amounting to 1 kg of food provided twice a day. Post experimental procedures, all animals were monitored 2-3 times a day to assess their responsiveness, urine and fecal output, mucous membrane color, water and food intake, pain assessment, and respiratory noise or effort which dictated humane endpoints if applicable.

#### 5.5.2 In Vivo Experimental Design, Surgical Procedure, and Assessment

An extended vertical incision was made at the midline on the ventral surface of the neck to expose the cervical trachea. Strap muscles and thyroid isthmus were retracted to allow the trachea to be elevated. A 4 cm x 2 cm tracheal defect was created over the anterior surface of the trachea. The removed tracheal segment was manipulated according to the following four groups prior to defect closure: 1) no manipulation (control) (*N*=3); 2) mucosa stripped (*N*=3); 3) mucosa stripped and replaced with bare SF-CVM graft (*N*=3); and 4) mucosa stripped and replaced with hiPSC-derived SF-CVM airway epithelial graft (*N*=4). For Group 4, prior to surgery, hiPSC-derived SF-CVM grafts were submerged and stained with 125 μM CellTracker™ 7-amino-4-chloromethylcoumarin (CMAC) Dye (Thermo Fisher) diluted in complete PneumaCult-ALI^TM^ media for 45 mins at 37°C, rinsed with DPBS. For Groups 3 and 4, SF-CVM grafts were attached to the removed segment via Fibrin Glue (Baxter International) and a running 4-0 PROLENE® polypropylene suture (Ethicon). The removed tracheal segment was re-attached using 4-0 PROLENE® polypropylene sutures to close the defect with the grafts oriented towards the luminal surface. A Valsalva maneuver was performed to check for and prevent any leaks during closure of the tracheal defect. Subsequently, Fibrin Glue (Baxter International) was applied to further seal the defect on the exterior of the trachea. The sternocleidomastoid muscle was subsequently mobilized and wrapped around the tracheal defect via 3-0 Vicryl® polyglactin sutures (Covidien). The animals were monitored daily for well-being and clinical symptoms of respiratory distress. On post-operative day (POD) 3, the wound was opened and a 6 cm circumferential portion of the trachea proximal and distal to the graft was resected for evaluation. The tracheal grafts were examined macroscopically, then segmented for separate assessment of histology (hematoxylin and eosin staining) and cell viability. For the latter, the grafts were stained with calcein-AM and ethidium homodimer-1 (according to kit instructions; Thermo Fisher) to assess cell viability and death, respectively. These samples were imaged immediately on a Zeiss LSM 710 NLO two-photon/confocal microscope. Post-operative area coverage of CMAC-stained cells was quantified using ImageJ software (NIH) by converting images into binary images via thresholding and analyzing pixel area.

### 5.6 Flow Cytometry

Efficiency of definitive endoderm induction on day 3 of differentiation was determined by assessing co-expression of c-KIT-APC (Thermo Fisher) and CXCR4-PE-Cy7 (Becton Dickinson) surface markers (*N*=3 for BU3NG hiPSC line and 110 hiPSC line). Intracellular staining for NKX2.1 and P63 was performed using a Cytofix/Cytoperm™ kit (Becton Dickinson). BU3NG Day 15 lung progenitors (*N*=3) were stained with Anti-TTF1 primary antibody [EP1584Y] (Abcam) and Alexa Fluor 647 secondary antibody (Life Technologies). Lung progenitors derived from hiPSC 110 were identified based on CD47 (high) and CD26 (low) expression via staining with Anti-human CD47 PerCP/Cy5.5 and Anti-human CD26 PE conjugate antibodies (Biolegend). Day 41-43 airway progenitors (*N*=3) were stained with rabbit monoclonal P63-PE antibody (Cell Signaling Technology). Samples were acquired on a LSRII SC flow cytometer (Becton Dickinson) and analyzed using FlowJo software (Becton Dickinson).

### 5.7 Immunocytochemistry

Post *in vitro* or *ex vivo* ALI culture, SF-CVM samples were fixed with 4% paraformaldehyde (Sigma Aldrich) for 10 minutes at room temperature (RT), permeabilized with 0.1% Triton X-100 (Sigma Aldrich) for 20 minutes at RT and blocked with 10% normal goat serum (Vector Laboratories) for 30 minutes at RT. Samples were stained with mouse monoclonal acetylated α-tubulin (Sigma Aldrich), mouse monoclonal mucin 5AC (Abcam), and rabbit polyclonal ZO-1 (Thermo Fisher Scientific) primary antibodies at 4°C overnight, and correspondingly stained with goat anti-mouse Alexa Fluor 546, goat anti-mouse Alexa Fluor 488, and goat anti-mouse Alexa Fluor 633 secondary antibodies (Life Technologies) for 1 hour at RT. All samples were counterstained with Hoechst (Sigma-Aldrich) and mounted. Three images per sample were captured on a Zeiss LSM 710 NLO confocal microscope and percentages of ciliated and goblet cells were manually quantified based on nuclear staining using ImageJ software (NIH).

### 5.8 Statistical Analysis

Analysis was conducted using a computer statistical software (GraphPad Software Inc., San Diego, CA, USA) with *P* values < 0.05 considered as significant. Descriptive statistics were presented as column bars with mean and standard error of the mean (SEM). Groups of two were analyzed with the unpaired two-tailed Student’s t-test and groups of more than two were analyzed using unpaired One-way ANOVA with Tukey’s post-hoc test. Data sets with more than two independent variables were compared using ordinary two-way ANOVA.

## Supporting Information

Supporting Information is available from Wiley Online Library or from the author.

## Acknowledgements

The authors are grateful to Dr. Darrell Kotton and Dr. Andras Nagy for providing the BU3NG and 110 hiPSC lines, respectively. The authors immensely thank Dr. Fabio Gava Aoki for his important contributions in designing the bioreactor air-liquid interface culture system. Gratitude is expressed to Dr. Cristina Amon and Dr. David Romero for their valuable inputs for the bioreactor design. Funding: R.V. was funded by the Henry White Kinnear Fellowship. This project was further funded by the University of Toronto Medicine by Design Initiative, funded by the Canada First Research Excellence Fund (CFREF; C1TPA-2016-18).

## Author Contributions

R.V. contributed to designing the overall study, designed and performed experiments, assisted in pig surgeries, analyzed data, and wrote the manuscript. A. E. M-A. developed the bioreactor air-liquid interface culture system, maintained bioreactor experiments, assisted in pig surgeries, and performed histology. S.R. set up and assisted in pig surgeries and performed histology. S.H. designed and performed the pig surgeries. T.K.W, G.K., and S.H. were principal investigators, supervised the overall work, and edited the manuscript.

## Data availability statement

The data that support the findings of this study are available from the corresponding author upon reasonable request.

## Table of Contents

Airway grafts derived from human induced pluripotent stem cells (hiPSC) and a Silk Fibroin-Collagen Vitrigel Membrane (SF-CVM) composite biomaterial are engineered and their ability to repair airway defects is demonstrated in a porcine model. The grafts successfully induce physiological proportions of mature epithelium. Importantly, hiPSC-derived SF-CVM grafts integrate with surrounding tissue, preserve airway patency, and maintain a viable, pseudostratified epithelium.

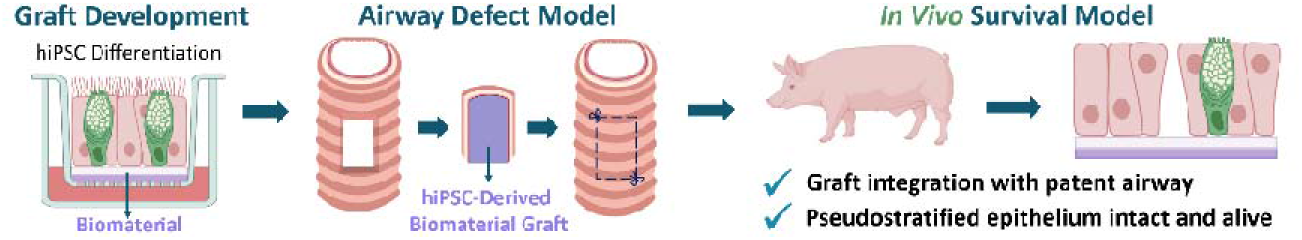

## Supporting Information

### Experimental Section/ Methods

#### HTEC Maintenance and ALI Culture

Normal human tracheal epithelial cells (HTECs) were collected from fresh tracheobronchial samples that were discarded during lung transplant operations using a previously described protocol.^[48]^ HTECs were maintained up to two passages in Basal Epithelium Growth Media (BEGM; Lonza) on Collagen I-coated (62 μg mL^−1^ PureCol, Advanced BioMatrix) tissue culture flasks. Passage 2 HTECs were seeded at a density of 100,000 cells (cm^2^)^−1^ on 24-well transwell inserts (VWR International) either coated with Collagen-I (control) or lined with SF-CVM. HTECs (*N*=3) were grown for 48 hours in submerged culture, with BEGM on both the apical and basal sides of the transwell insert. They subsequently underwent an “air-lift” and were nourished only from the basal side with Bronchial Air Liquid Interface (B-ALI; Lonza) up to 31 days. SF-CVM was glued onto the transwells using Fibrin Glue (TISSEAL Fibrin Sealant; Baxter International).

#### ALI Optimization Study on SF-CVM

BU3NG hiPSC-APs were seeded on 24-well transwell inserts lined with SF-CVM at a density of 500,000 cells (cm^2^)^−1^ and allowed to form a monolayer in submerged culture for 5 days in complete PneumaCult-ALI^TM^ (Stem Cell Technologies) media (containing dexamethasone instead of hydrocortisone, 1% Antibiotic-Antimycotic (Thermo Fisher), and 1% gentamicin (Wisent)) with further supplementation of 2 μм Dorsomorphin and 10 μм SB431542. On day 5, air-liquid interface (ALI) culture was begun by feeding only the bottom chamber with complete PneumaCult-ALI^TM^ media for an additional 31 days (day 72-74). During this ALI culture phase, SF-CVM grafts either received no supplementation, or one of the following 12 treatments: 10 ng basic FGF (bFGF), 100 ng bFGF, 5 μм DAPT, 10 μм DAPT, 100 nM DBZ, 1000 nм DBZ, 100 ng mL^−1^ dickkopf-1 (DKK-1), 200 ng mL^−1^ DKK-1, 10 ng mL^−1^ insulin growth factor-1 (IGF-1), 100 ng mL^−1^ IGF-1, 10 ng mL^−1^ interleukin-6 (IL-6), and 20 ng mL^−1^ IL-6 (*N*=3 for all groups). The resulting grafts were assessed for terminal differentiation into ciliated and goblet cells via immunocytochemistry. Three images per sample were captured on a Zeiss LSM 710 NLO confocal microscope and percentages of ciliated and goblet cells were manually quantified based on nuclear staining using ImageJ software (NIH).

**Figure S1.**
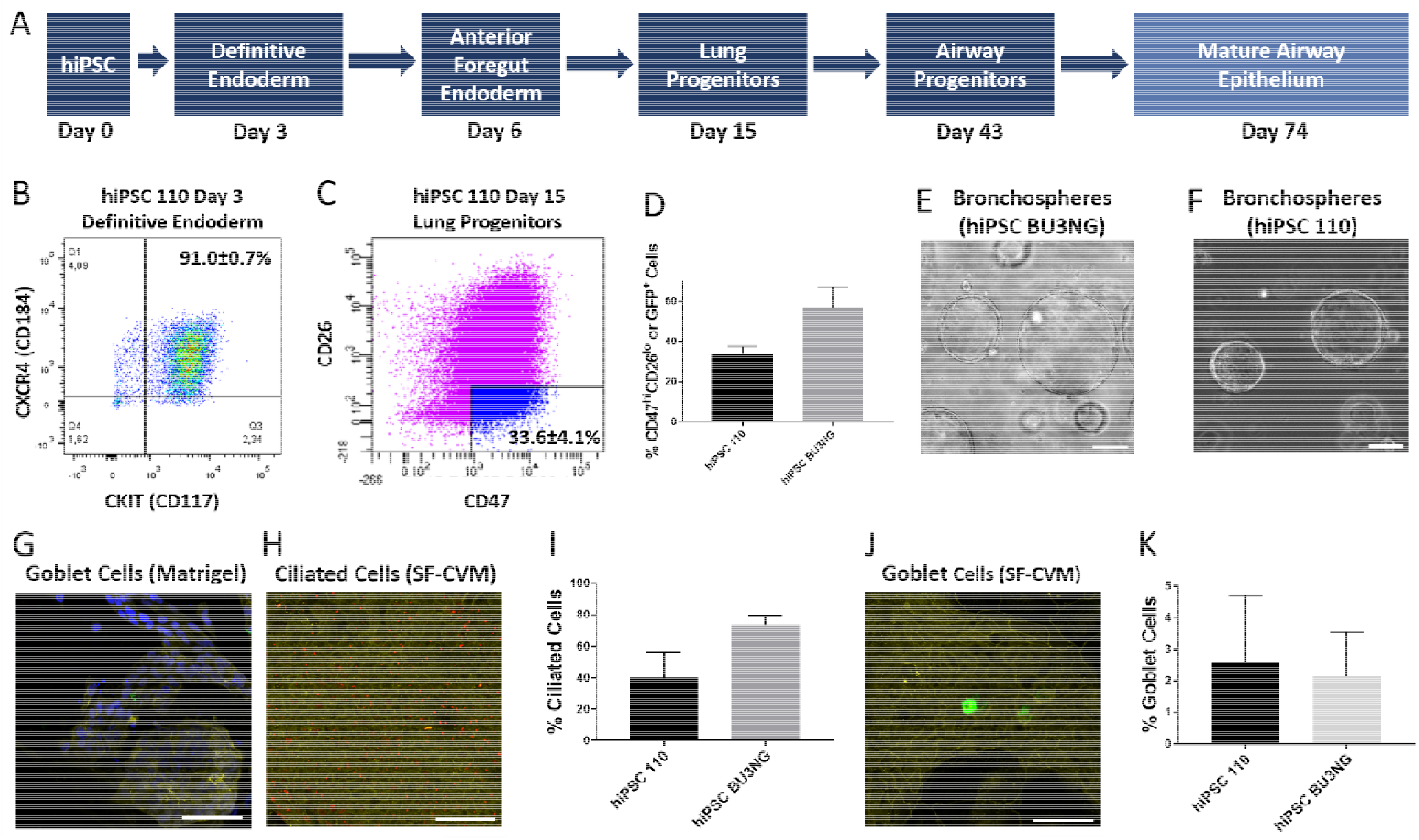
Mucociliary SF-CVM airway epithelial grafts derived via directed differentiation of the 110 hiPSC cell line. Directed differentiation schematic for achieving mucociliary airway epithelium (**A**). Representative flow cytometry plots of CKIT^+^CXCR4^+^ cells (*N*=3) on Day 3 (**B**) and CD47^hi^CD26^lo^ cells (*N*=4) on Day 15 (**C**). Comparison of lung progenitors produced by the 110 and BU3NG hiPSC lines (**D**; *N*=4; *P*=0.10). Sorted lung progenitors of both the BU3NG (**E**) and 110 (**F**) hiPSC lines were embedded in Matrigel at 1000 cells μL^−1^ and 3D cultured to generate bronchospheres by Day 43. Dissociated Day 43 cells were seeded on Matrigel-coated or SF-CVM-laden transwells at 500,000 cells (cm^2^)^−1^ and air-lifted on day 5 to initiate ALI culture. Samples were fixed on day 31 of ALI (Day 74), stained using antibodies against acetylated α-tubulin (red; ciliated cells), zonula occludens-1 (yellow; tight junctions), and mucin 5AC (green; goblet cells), and counterstained with Hoechst (not shown). Patchy epithelial layer on Matrigel-coated transwells (**G**). Representative images of ciliated (**H**) and goblet (**J**) cells on SF-CVM. No significant differences were observed amongst percentages of ciliated cells (*P*=0.12; *N*=3; **I**) and goblet cells (*P*=0.87; *N*=3; **K**) between the 110 and BU3NG hiPSC lines on Day 74 according to two-tailed Student’s t-test. Scale bars: 50 μm. Mean ± SEM.

**Figure S2.**
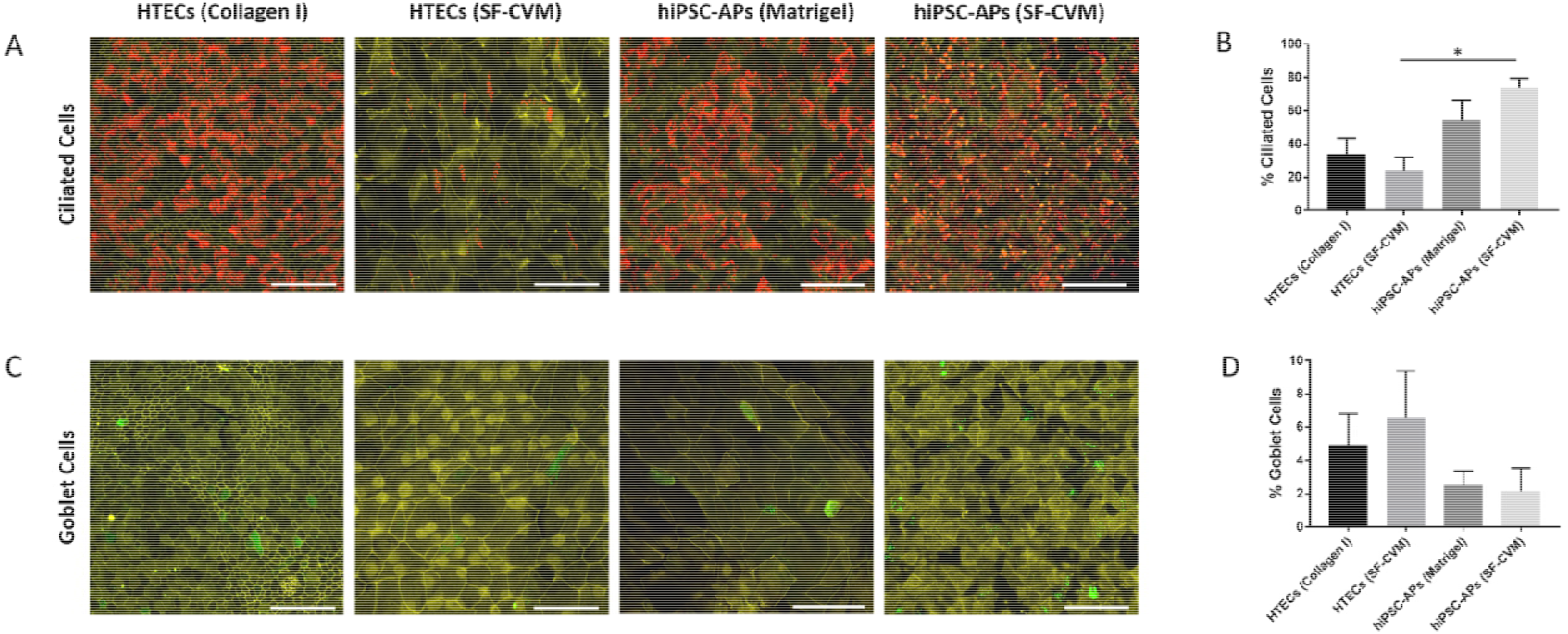
hiPSC-APs differentiate into significantly higher ciliated cell populations on SF-CVM compared to HTECs. HTECs seeded on Collagen I-coated or SF-CVM-laden transwells at a density of 100,000 cells (cm^2^)^−1^ and ALI culture within 48 hours. BU3NG hiPSC-APs were seeded on Matrigel-coated or SF-CVM-laden transwells at 500,000 cells (cm^2^)^−1^ and air-lifted on day 5 to initiate ALI culture. Samples were fixed on Day 31 of ALI, stained using antibodies against acetylated α-tubulin (red; ciliated cells), zonula occludens-1 (yellow; tight junctions), and mucin 5AC (green; goblet cells), and counterstained with Hoechst (not shown). Representative images of resulting HTEC or hiPSC-AP-derived ciliated (**A**) and goblet (**C**) cells. Significantly higher ciliated cell populations (*P*<0.05) generated by hiPSC-APs compared to HTECs on SF-CVM according to One-way ANOVA and Tukey’s post-hoc tests (**B**). Quantified goblet cell populations without significant differences amongst groups (**D**). Scale bars: 50 μm. *N*=3; Mean ± SEM.

**Figure S3.**
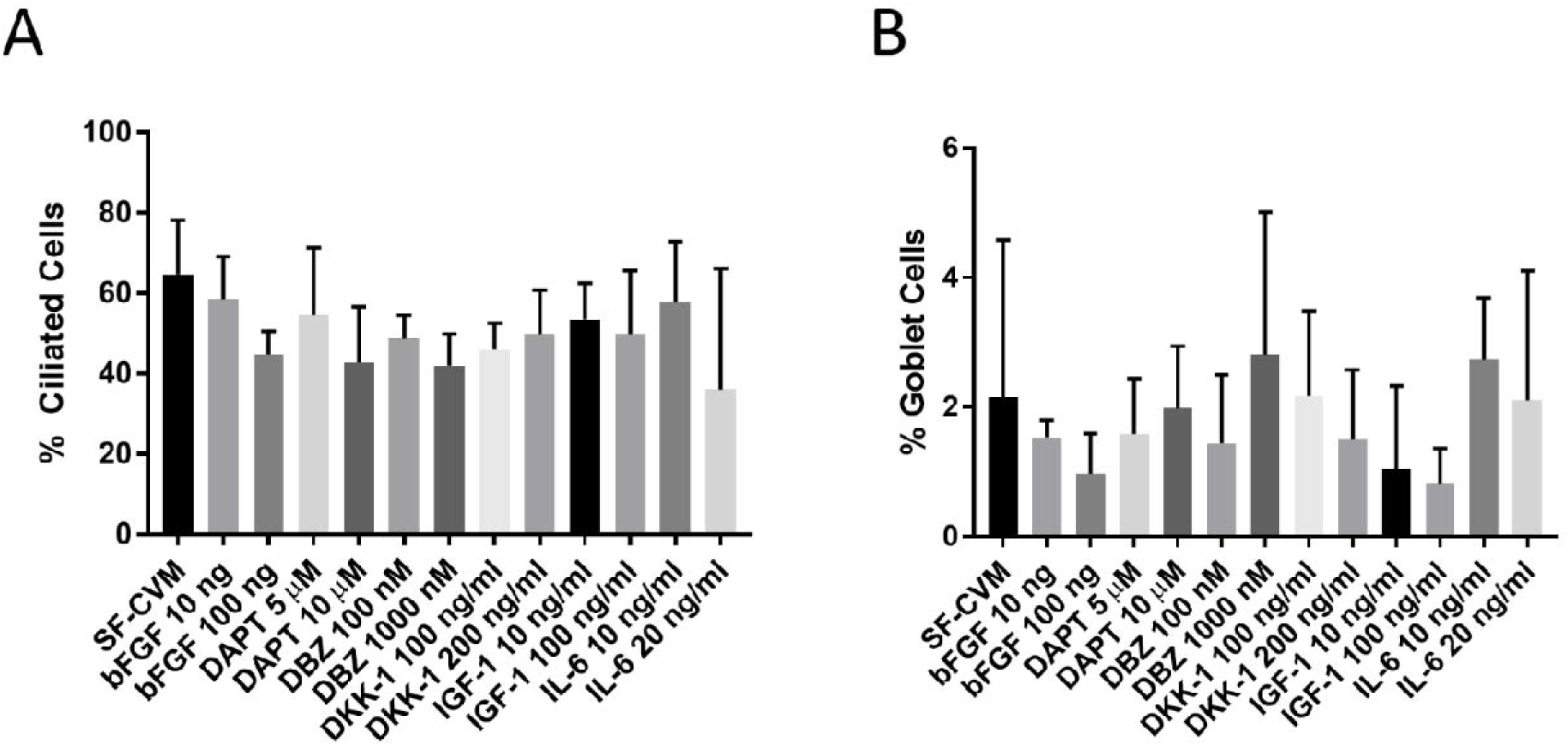
Media supplementation during ALI culture does not affect ciliated or goblet cell differentiation on SF-CVM. BU3NG hiPSC-APs were seeded on SF-CVM-laden transwells at 500,000 cells (cm^2^)^−1^ and air-lifted on day 5 to initiate ALI culture for an additional 31 days. SF-CVM grafts were cultured in ALI media with or without supplementation with bFGF, DAPT, DBZ, DKK-1, IGF-1, or IL-6. Analysis of quantified ciliated (**A**) and goblet (**B**) cells found no significant differences amongst groups via One-way ANOVA and Tukey’s post-hoc tests. *N*=3; Mean ± SEM.

**Figure S4.**
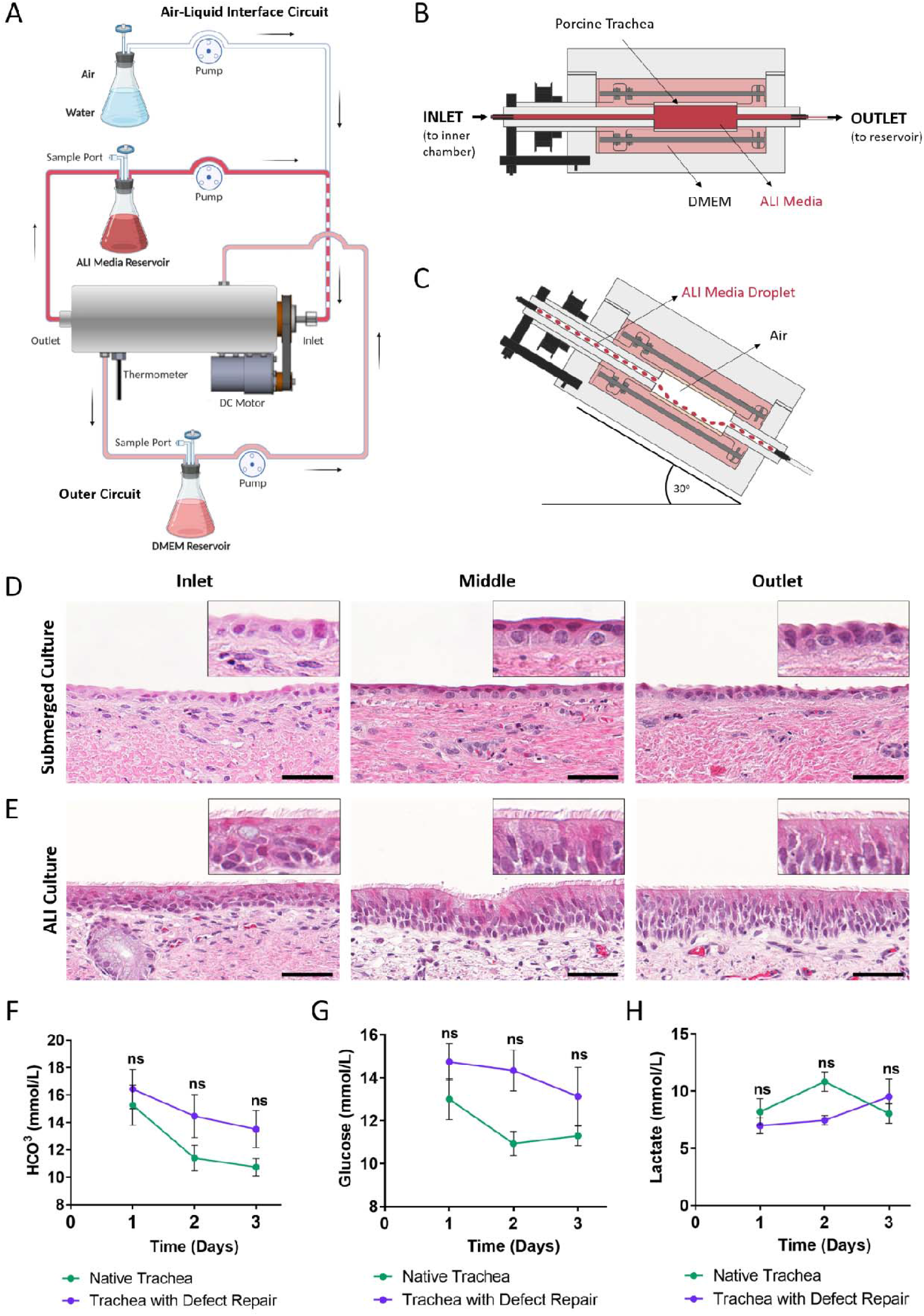
Bioreactor ALI culture system maintains pseudostratified, ciliated epithelium of porcine tracheae across 7 days of culture. Schematic of the bioreactor ALI culture circuit (partially created with BioRender.com) (**A**). Bioreactor setup diagrams for submerged (**B**) and ALI (**C**) culture. Native porcine tracheal segments from the inlet, middle, outlet regions of the bioreactor stained with hematoxylin and eosin after 7 days of submerged (**D**) or ALI culture (**E**). Epithelial cell metabolism as evidenced by bicarbonate (**F**), glucose (**G**), and lactate (**H**) quantities for both native and repaired tracheae across 3 days of ex vivo ALI culture with no statistical differences found. Statistical analysis of metabolite concentrations performed using Two-way ANOVA. *N*=3; Scale bars: 50 μm.

